# Differences in Cajal-Retzius cell density and postnatal persistence across cortical areas revealed by a novel intersectional genetic labeling approach

**DOI:** 10.1101/2025.04.09.647949

**Authors:** Kristian Moan, Robert Machold, Giulia Quattrocolo

## Abstract

Cajal-Retzius (CR) cells are glutamatergic neurons that transiently populate the most superficial layer of the isocortex and allocortex during development, serving an essential role during both prenatal and early postnatal brain development. Notably, these cells disappear from most cortical areas by postnatal day 14, but persist for much longer in the hippocampus. We developed a novel intersectional genetic labeling approach for CR cells that captures almost all of the TRP73-positive CR cells throughout the isocortex and allocortex. This intersectional strategy offers several advantages over previous methods commonly used for CR cell targeting. Here, we applied this new CR cell labeling strategy to investigate the distribution and persistence of CR cells throughout the whole mouse brain, at four different postnatal ages. We observed that the initial CR cell density and the rate of their disappearance varies substantially across different brain areas during development. Strikingly, we observed variation in cell death rate even between adjacent cortical subregions: comparing the medial and the lateral entorhinal cortex, the former retains a high density of CR cells for several months in contrast to the latter. Our results present a necessary revision of the phenomenon of CR cell persistence, showing that, in addition to hippocampus, several other cortical areas maintain a high density of these cells beyond the first two postnatal weeks.

## INTRODUCTION

Cajal-Retzius (CR) cells are a transient glutamatergic neuronal population that plays a critical role in early cortical development. During embryogenesis, CR cells, together with subplate neurons, form the cortical preplate (Meyer et al., 1999). Once positioned in the marginal zone, cortical CR cells guide the inside-out radial migration of newly born pyramidal neurons (PNs) to their correct lamina, in part through secretion of the reelin protein (Frotscher, 1998; Ogawa et al., 1995). This regulation of PN migration by CR cells is required for the formation of the six-layered isocortex (D’Arcangelo et al., 1995; Ogawa et al., 1995). CR cells are found throughout the allocortex, spanning the lateral olfactory tract, piriform cortex, and the parahippocampal cortex (Causeret et al., 2021; de Frutos et al., 2016; Elorriaga et al., 2023). In the hippocampus, CR cells have been proposed as transitional targets for incoming axons reaching the hippocampus (Ceranik et al., 2000). Beyond their roles in guiding cell migration and axonal pathfinding, CR cells are crucial for the development of the local circuits of both the isocortex (Damilou et al., 2024; Riva et al., 2023) and the hippocampus (Glærum et al., 2024).

After birth, as development progresses, the transient nature of CR cells becomes apparent. In the mouse, after about postnatal day 14 (P14), CR cell numbers are greatly reduced (Chowdhury et al., 2010; del Río et al., 1995) through programmed cell death (Ledonne et al., 2016). Intriguingly, the hippocampal CR population does not decrease at the same rate as their isocortical equivalent (Del Río et al., 1996). This trend continues throughout development, such that by the beginning of adult stages, virtually all CR cells are gone, except for the remaining hippocampal population (Anstötz et al., 2018a; Del Río et al., 1996; del Río et al., 1995; Super et al., 1998). Given this unambiguous difference, it has been speculated that hippocampal CR cells might perform additional functions in postnatal development, or that their function is needed for a longer period (Anstötz et al., 2018b). The early arrival of these cells in cortical regions, their importance during development, and their differential persistence across isocortical and hippocampal areas has made CR cells a prime focus of neurodevelopmental studies.

There is a long history of identifying and studying CR cells. Since their initial recognition by Ramon y Cajal in 1890 (Gil et al., 2014; Ramón y Cajal, 1890; Retzius, 1893), an increasing repertoire of characteristics has been established to help identify and manipulate these cells. First, CR cells have a unique bipolar morphology and are exclusively localized in the most superficial layer. Second, a number of molecular markers have been identified to distinguish these cells. In particular, reelin is recognized as the most comprehensive marker, being expressed in all CR cells (Bielle et al., 2005); however, this protein is also expressed by GABAergic interneurons present in the same layers (Pelkey et al., 2017). TRP73 (p73) is a commonly used marker that is very specific for CR cells (Elorriaga et al., 2023), but it does not label CR cells originating from the pallial-subpallial border (PSB) (Griveau et al., 2010). Historically, Calretinin has been widely used to identify CR cells, but has proven to be somewhat unreliable (Causeret et al., 2021; Weisenhorn et al., 1994). In addition, Calretinin is also expressed by a subset of neocortical interneurons (Barinka & Druga, 2010). Ultimately, the experimental characterization and manipulation of a specific cell type requires transgenic animal models or viral vectors to target it. Several transgenic mouse lines, each with different degrees of specificity, have been employed or developed for the purpose of manipulating CR cells (Pde1c-Cre: (Osheroff & Hatten, 2009) ΔNp73Cre: (Tissir et al., 2009; Yoshida et al., 2006); Frizzled10-CreERT2 (Gu et al., 2009); CXCR4-EGFP (Marchionni et al., 2010), but their characterization especially in the hippocampus and at adult stages has been incomplete to date.

Here, we describe and characterize a novel intersectional genetic strategy to target CR cells that leverages the observation that CR cells express neuron-derived neurotrophic factor (NDNF) (Kuang et al., 2010). We combined a recently developed NDNF driver line (NDNF-flox-FlpO; (Chamberland et al., 2024)) with a Pde1c-Cre line (Osheroff & Hatten, 2009) and the intersectional tdTomato reporter Ai65 (Jax #021875; (Madisen et al., 2015)) to label CR cells. This cross (CR:tdTomato) exhibited an impressive specificity for CR cells throughout early postnatal stages and up to adulthood, and allowed us to investigate the difference in CR cell density, distribution, and persistence across the development of the isocortices and the allocortex. Thus, this genetic targeting approach represents an important experimental tool for studying CR cells across both isocortex and allocortex. Our data indicate that the commonly held view that CR cells uniformly disappear from all isocortical regions within the first two weeks of postnatal development will need to be revised. Indeed, we demonstrate that in addition to the hippocampus, several cortical areas have a prolonged persistence of CR cells beyond P14, pointing to a wider function for these cells in postnatal maturation of brain circuitry.

## MATERIALS AND METHODS

### Animals

All experiments were conducted in compliance with protocols approved by the Norwegian Food Safety Authorities and European Directive 2010/63/EU (FOTS ID 24847 and 31016). Pups were separated from the mother at postnatal day 21. Adult animals were housed with up to five animals per cage and kept in an inverted 12-hour light/dark cycle, with enriched cages, and food and water ad libitum.

Pde1c-Cre transgenic mice (B6.FVB(Cg)-Tg(Pde1c-Cre)IT146Gsat/Mmucd, MMRRC 030708) (Osheroff & Hatten, 2009) were bred with NDNF-flox-FlpO transgenic mice to create compound heterozygous *Pde1c-Cre^+/−^;NDNF-flox-FlpO^+/−^* mice. This line was subsequently bred with homozygous Ai65 (B6.Cg-Gt(ROSA)26Sortm65.2(CAG-tdTomato)Hze/J) (21; JAX stock #021875) animals to produce *Pde1c-Cre^+/−^;NDNF-flox-FlpO^+/−^*;Ai65*^+/+^* experimental animals (CR:tdTomato). Animals of both sexes were used for the experiments.

### Perfusion and section preparation

Animals were first anesthetized with 0.1 ml isoflurane in a closed chamber and subsequently injected with a lethal dose of pentobarbital (100mg/kg, 0.1ml, IP, SANVIO Pharma AS). Following, the animal was transcardially perfused with ice cold (4 °C) PBS for 5 min (3.5ml / min) followed by freshly made, ice cold, paraformaldehyde (PFA, pH 7.4, Merck Life Sciences AS) for 5 min (3.5ml / min). After recovery of the brain, it was post-fixated in PFA for 16-24h in the fridge. Brains were cryoprotected in increasing concentrations of sucrose (15%, 30%) in preparation for sectioning. P15 – P90 brains were cut in either sagittal or coronal orientation, on a freezing microtome (Sliding microtome HM 430, Thermo Scientific, Walldorf, Germany) with 50 µm thickness, in three series, before being stored in antifreeze (40% 125 mM phosphate buffer (PBS), 30% glycerol, 30 % ethylene glycol). P1 brains were first embedded in OCT mounting media (VWR international), before being cut either sagittal or coronally, on a cryostat (Cryostar NX70, Thermo Scientific, Walldorf, Germany) with a 30 µm thickness, in three series, immediately mounted on microscope glass slides (Thermo scientific, Superfrost Plus, 25×75x1 mm) and stored at −20 °C.

### Immunohistochemistry

#### P15 to P90 brains

One series of sections from each brain were used for whole brain/hemisphere immunolabeling. Floating sections were initially washed in a 12 well-dish, in PBS 3 × 10 min, followed by a blocking step containing 10% normal donkey serum (Jackson ImmunoResearch) and 0.1% Triton X-100 (Merk) dissolved in PBS, for 30 min. Next, sections were incubated in primary antibodies in a buffer containing 1% normal donkey serum and 0.1% Triton X-100 dissolved in PBS, for three overnights at 4 °C, while rotating. Primary antibodies: Guinea pig anti-NeuN 1:1000 (Merk, ABN90P), Goat anti-mouse reelin 1:500 (Biotechne RandD systems, AF3820), Rabbit anti-P73 1:1000 (Abcam, ab40658), Rat anti-RFP 1:1000 (ChromoTek, AB_2336064). After primary antibody incubation, sections were again washed in PBS 3 × 10 min, before incubation in secondary antibodies, in similar buffer as primary antibodies, overnight at room temperature while rotating. Secondary antibodies: Donkey anti-Guinea pig AF 405 1:250 (Jackson Immuno Research Lab, 706475148), Donkey anti-Goat AF 488 1:500 (Invitrogen A32814), Donkey anti-Rat AF 594 1:500 (Invitrogen, A21209), Donkey anti-Rabbit AF 647 1:500 (Invitrogen, A31573). Lastly, the sections were washed in PBS 3 × 10 min, before being mounted on microscope glass slides (Thermo Scientific, Superfrost Plus®, 25×75x1) and dried on a heating pad at ∼40 °C. After drying, whole microscope glasses were washed in deionized water (dH_2_O), before being dried again. The glasses were coverslipped (Menzel-Gläser, 24×50 mm, #1) using Fluoromount (Invitrogen, Fluoromount-G) as embedding medium.

#### P1 brains

One series of P1 brain sections was used for whole brain/hemisphere immunolabeling. All washings and incubations were done on mounted sections, on glass. To prepare the glass for washing, a circle was drawn on the glass, around the mounted sections, with a PAP pen (SuperHT, pap pen). Initially, sections were washed, in PBS 3 × 10 min, followed by a blocking step containing 10% normal donkey serum (Jackson ImmunoResearch,) and 0.1% Triton X-100 (Merk) dissolved in PBS, for 1 hour. Next, sections were incubated in primary antibodies in a buffer containing 1% normal donkey serum and 0.1% Triton X-100 dissolved in PBS, for one overnight at 4 °C, while rotating. After primary antibody incubation, sections were again washed in PBS 3 × 10 min, before incubation in secondary antibodies, in similar buffer as primary antibodies, 2 hours at room temperature, while rotating. All antibodies were the same as described for P15 to P90 brains. Lastly, the sections were washed in PBS 3 × 10 min, followed by 10 min with dH_2_O. The glass was then coverslipped while the sections were still wet, with the same cover glass and mounting media as mentioned above.

### Imaging

#### Confocal imaging

Specific brain regions were imaged at high-resolution using a Zeiss LSM 880 Indimo confocal module attached to a Axio-ImagerZ.2 (Carl Zeiss, Jena, Germany). All confocal images were acquired with a Plan-Apochromat 20x/0.8 M27 objective, with a resolution of ∼0.26 µm in X and Y, and ∼1.0 µm in Z. (pinhole size of ∼1 Airy Unit, 8-bit depth, ∼0.5 μsec pixel dwell, an averaging of 1 and unidirectional scan direction). Four channels were acquired, AF 405 was excited by 405 nm laser, with emission filter set at 410-483 nm. AF 488 was excited by an Argon 488 nm laser, with emission filter set at 491-553 nm. AF 594 was excited by diode-pumped solid state (DPSS) 561 nm laser, with emission filter set at 588-660 nm. Lastly, AF 647 was excited by Helium Neon (HeNe 633) red 633 nm laser, with emission filter set at 660-755 nm. All images were acquired as Z-stacks and merged as a maximum intensity projection (MIP). Images were saved as .czi format and the confocal was controlled with Zen black v.2.1 SP3 (RRID:SCR_013672).

#### Scanner imaging

Full series of mounted sections were imaged with a Zeiss AxioscannerZ.1 (Carl Zeiss) controlled with Zen Blue software. All scanner images were acquired with a Plan-Apochromat 20x/0.8 M27 objective, with a resolution of ∼0.33 µm in X and Y, and 4.0 µm in Z. All images were acquired as Z-stacks with four slices across ∼15 µm. Four channels were acquired, AF 405 was excited by led-module 365 nm, with emission filter set at 412-438 nm. AF 488 was excited by led-module 470 nm, with emission filter set at 501-538 nm. AF 594 was excited by led-module white with beam splitter 560, with emission filter set at 570-640. Lastly, AF 647 was excited by led-module 625 nm, with emission filter set at 662-756 nm. All images were saved as .czi format with one image file containing the full microscope glass. The images were later split into single sections.

### Cell counting

#### Pre-processing

Before starting cell counting, the scanner images were pre-processed with several custom-made scripts, using Fiji/ImageJ (V1.154d, National Institute of Health, USA, https://imagej.net/). After being split into single files from the scanner images, the resulting images were merged into a MIP and saved in .tif format. Next, the different channels in the images were split into separate images, saving the channel for NeuN in one group and the channel for tdTomato in another group. Additionally, images were compressed by saving in .png format. The NeuN group went through a final process whereby the images were brightness and contrast adjusted for easier user-guided atlas alignment.

#### Training the counter

A subset of images was chosen for training. Five sagittal sections, distributed from most lateral to most medial, from one brain per age group, except for P1, were used to train the counting software. Ilastik v.1.4.0 ((Berg et al., 2019); RRID:SCR_015246) was used to do all automatic counting. First, patterns for pixel classification and object segmentation were trained with a user guided module to a quality that could catch as many cells as possible while still avoiding the inclusion of incorrect signals. For the P1 dataset, a subset of images (14), spanning from most lateral to medial and anterior to posterior, was chosen for training, using the same software. The following training steps were done in the same fashion as mentioned before.

#### Cell counting

The QUINT ((Yates et al., 2019); RRID:SCR_016854) workflow, with the Allen mouse brain atlas common coordinate framework version 3 (CCFv3) ((Wang et al., 2020); RRID:JCR_020999; RRID:JRC_021000) was used to do automatic cell counting and atlas registration. All images were cell counted with the trained software, first with pixel classification. Subsequently, one pixel classification file per images was saved in .H5 format. Next, the resulting pixel classification files were used to do object classification, and the resulting files were saved in .H5 format. Lastly, the final .H5 files were converted to segmentation bitmaps using ImageJ and adding the “Glasbey-inverted” look-up table. These bitmaps were saved as .png and named with the animal ID and numbered. All the segmented bitmaps were visually examined to look for obvious errors in the counting. In instances where the counter had classified artifacts as cell signals, the particular erroneous segmentation was masked using Adobe Photoshop (v.26.5.0, Adobe; RRID:SCR_014199).

#### Atlas registration

The NeuN images were aligned to a mouse brain atlas using the registration tool QuickNII v.3.2017 ((Puchades et al., 2019); RRID:SCR_016854) for P30-90 brains, or DeMBA v.2 beta for P1-15 brains (Carey et al., 2024). QuickNII filebuilder software was first used to make a read-file of the image folder. Next, the read-file was opened in QuickNII, and linear registration was initiated with the atlas for the respective age. The only exception was for the P1 group, where the P4 model was used as it was the youngest available model. After manual registration of every section to the corresponding atlas image, the alignment file was saved in .json format. This file was used to open the same project in the complementary non-linear registration tool VisuAlign ((Puchades et al., 2019); RRID:SCR_017978) v.2022 or v.DeMBA-CCFv3-2022 (Carey et al., 2024). Here, all atlas images were transformed non-linearly to precisely fit the brain section images. The resulting file acted as the atlas anchor file and was saved in .json format. The individual transformed atlas images were exported in .FLAT format.

#### Quantification of cell counts

As the last step, Nutil ((Groeneboom et al., 2020); RRID: SCR_017183) v.0.8.0 was used to quantify the cell counting by merging the transformed atlas images from the registration with the segmentation files from the automatic cell counter. The resulting files included specific counts from all areas defined from the CCFv3, and coordinates in this space, for all counted cells.

### Visualization

The heatmaps were created using the Brainglobe-heatmap v.0.2 (Claudi et al., 2023) software from the Brainglobe software python package with the Brainglobe atlas API ((Claudi et al., 2020); RRID:SCR_023848). All figures were made or assembled using Adobe Illustrator (v.29.4, Adobe; RRID:SCR_010279). For figure 1, the mouse icon was obtained from BioRender (www.biorender.com).

### Statistical analysis

All statistical analyses were conducted using GraphPad Prism version 10.0.0 for Windows (GraphPad Software, Boston, Massachusetts USA, www.graphpad.com). The density data was averaged between the three subjects before they were normalized. Density data used for heatmap figures were normalized in two different ways. For data normalized across all ages, we did a min-max normalization, assigning the highest density from across all ages as 1, and the lowest density as 0, while all other densities were assigned a value between 1 and 0, depending on their value. Similarly, for normalization over same age, we performed a min-max normalization only with density values from the same age.

## RESULTS

### Intersectional genetic targeting of CR cells

Previous work from our lab and others have utilized a Pde1c-Cre line to target CR cells at postnatal ages (Anstötz et al., 2018a; Glærum et al., 2024). However, this approach requires local viral injection with Cre-dependent AAV constructs and is impractical for brain-wide labeling as Pde1c is also expressed in non-CR populations to some extent, like a subset of layer 5 cortical PN (Tasic et al., 2016). To develop a genetic strategy to label CR cells efficiently and precisely across the whole mouse brain, we tested whether intersectional genetics with another CR cell-expressed gene could yield the desired result. Since the gene NDNF is also expressed in CR cells (Kuang et al., 2010), we took advantage of a recently developed NDNF-flox-FlpO driver line where FlpO is expressed from NDNF+ cells only following Cre-mediated removal of the last coding exon. As outlined in **Figure 1**, we generated Pde1c-Cre^+/−^;NDNF-flox-FlpO^+/−^ compound heterozygous mice and crossed these with homozygous Cre and Flp dependent tdTomato reporter mice (Ai65; *Rosa26tdtomatoAi65^+/+^*) to produce CR:tdTomato mice (Pde1c-Cre; NDNF-flox-FlpO; Ai65). We validated the accuracy and efficiency of this CR cell labeling approach and performed cell counting of all tdTomato labeled CR cells across whole mouse brains at four different developmental timepoints (P1, P15, P30, and P90) to investigate whether there are differences in CR cell density and persistence across the brain. To gain an overview of the expression patterns in our CR:tdTomato mouse, we analyzed whole brain sagittal sections from each age group and observed that labeled cells were selectively localized in the most superficial layer of the isocortex and the allocortex, and appeared to be CR cells based on their morphology (**Fig 2.A-B, Suppl.Fig 1. A**).

**Fig. 1.**
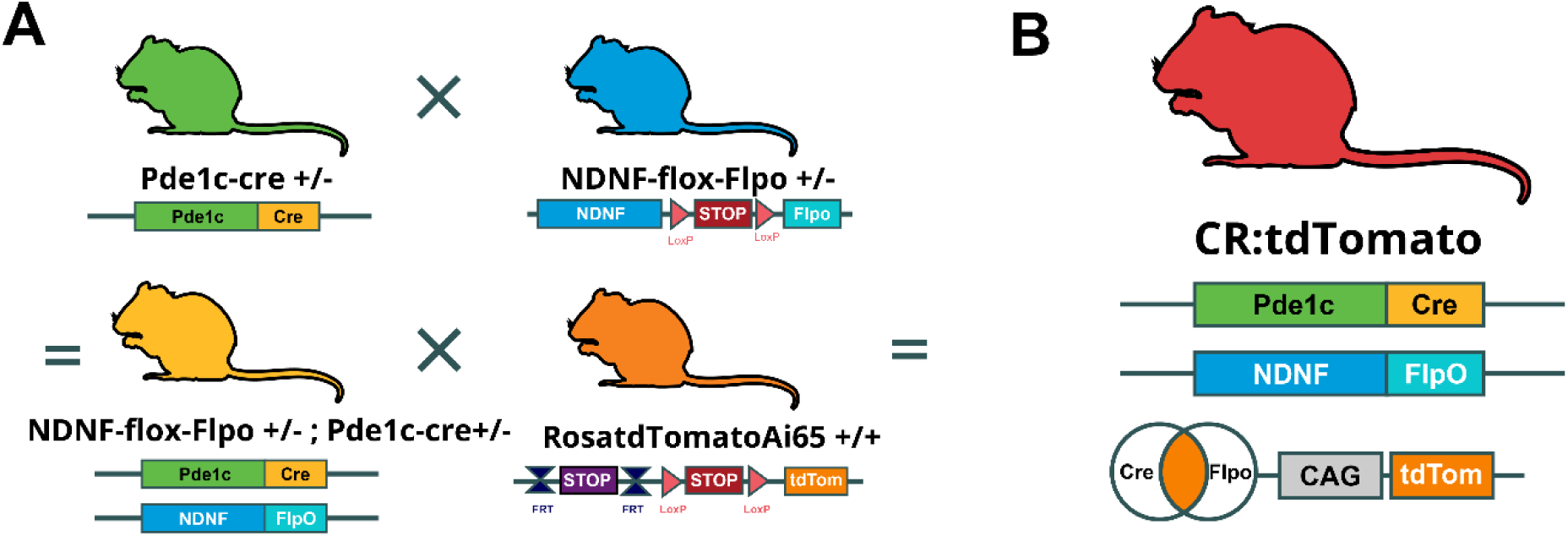
Transgenic strategy for labeling CR cells (CR:tdTomato). (A) All transgenic mouse lines used, with schematics under each respective mouse line depicting the relevant genes, recombinases, loxP-flanked sites, and stop cassette-regions. Pde1c-Cre mice were mated with NDNF-flox-FlpO mice to create NDNF-flox-FlpO; Pde1c-Cre compound heterozygous mice, which were then mated with homozygous RosatdTomatoAi65 reporter mice to create our CR:tdTomato mice (Pde1c-Cre; NDNF-flox-FlpO; Ai65). (B) Summary schematic of the CR:tdTomato mouse outlined in A. Venn diagram illustrates how cells expressing both recombinases will express the tdTomato fluorescent protein under the control of the Ai65 CAG promoter.

**Fig. 2.**
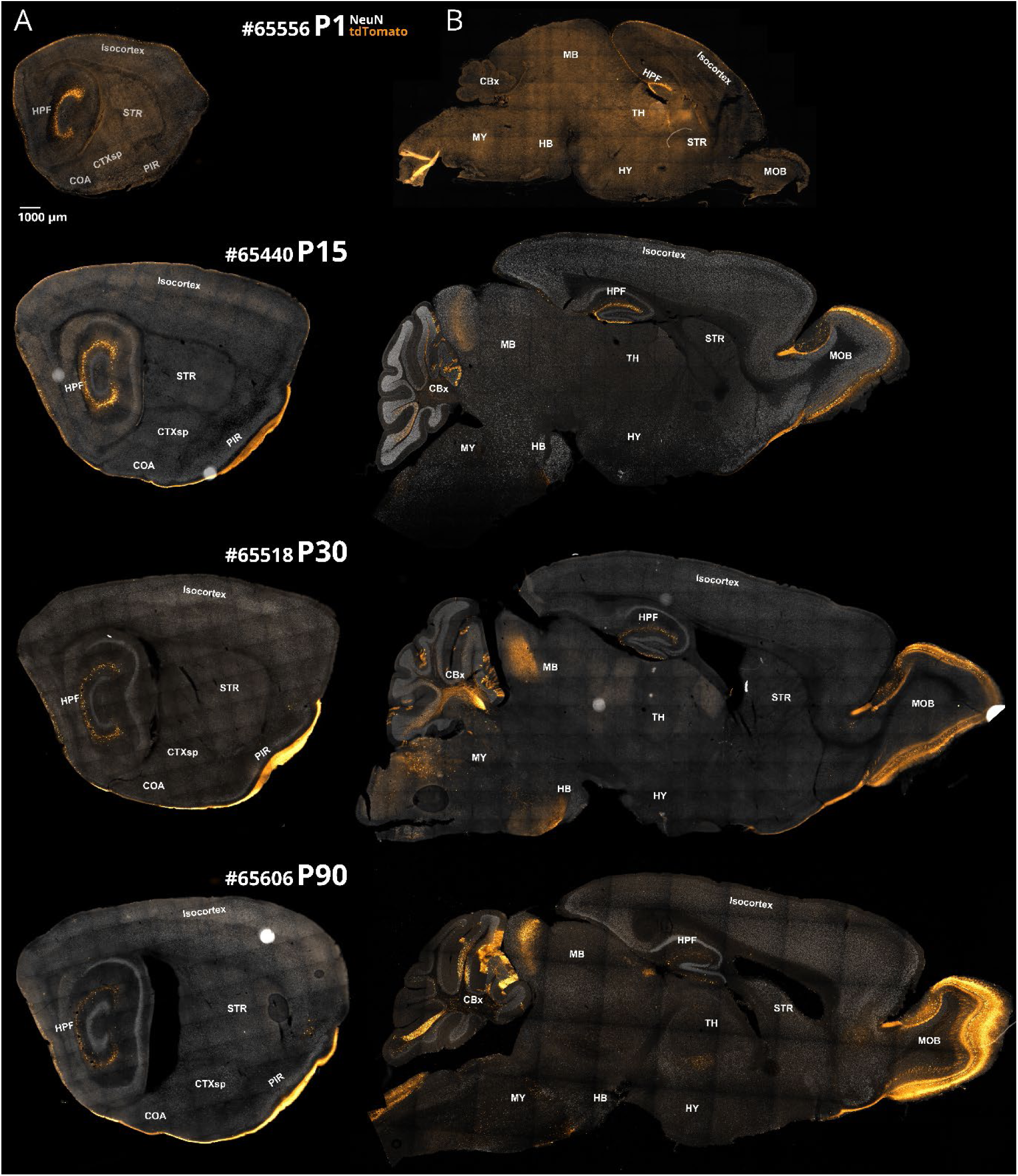
Cell labeling in the CR:tdTomato mouse line across four age points. (A) Sagittal sections from CR:tdTomato brains (P1, P15, P30 and P90) at a lateral position. (B) Sagittal sections from CR:tdTomato brains (P1, P15, P30 and P90) at a medial position. Primary areal subdivisions of the brain are indicated with Allen Brain Atlas abbreviations. Scale bar (top left) for all images shown represents 1 mm. Sections from P1 through P90 brains are arranged from top to bottom.

In addition to the labeling of putative CR cells in isocortical and allocortical areas, we also observed tdTomato expression in several non-CR populations in the brain. We noted a small population of non-CR neurons from P30 onward in the deep layer of the piriform cortex (**Suppl.Fig 1. D**). In addition, starting at P15, a thick bundle of tdTomato+ axonal fibers appeared, travelling ventrally through the piriform cortex, ultimately all the way to the entorhinal cortex (EC) (**Fig. 2A, P15**), most likely stemming from non-CR cells in the main olfactory bulb (MOB) (**Fig. 2B, P15, Suppl.Fig 1. C**). From P15, expression was observed in cerebellar Purkinje cells, identified based on layer location and their distinctive morphology (**Supp.Fig 1. B**). The Purkinje cells expressing tdTomato appeared to be situated in lobule II, III and IX, although not exclusively, as Purkinje cells were observed sparsely throughout most of the cerebellar lobules. In the olfactory bulb (OB), neurons in the mitral and outer plexiform layers expressed tdTomato from P15 onward, likely constituting the main excitatory neuronal populations in the OB, the mitral cell and tufted cells, respectively. The mitral cells are the most probable source of the axonal projections observed passing through piriform towards EC, as these cells are the main cortically projecting neuronal population in the OB (Imamura et al., 2020). Additionally, neurons in the association OB expressed tdTomato, and, from P30 onward, neurons in layer 2 anterior olfactory nucleus were also labeled. In the dorsal medulla, we observed several scattered populations of tdTomato+ neurons from P15 onward. These were located generally at the most dorsocaudal part and likely consist of neurons in the nucleus of the solitary tract (NTS), cuneate nucleus (CU), medullary reticular nucleus (MDRN), and spinal nucleus of the trigeminal (SPVC) (**Supp.Fig 1. C**). We inferred that all these non-CR neurons express both Pde1c and NDNF, but likely with different timing as the onset of fluorescence was not the same between these regions. Notwithstanding these additional labeled populations, our visualization highlighted the absence of any non-CR cells in all cortical areas, except deep layer piriform cortex. In addition, the decrease of CR cell numbers, and the difference in persistence between isocortex and hippocampal region, from P1 to the older ages, was clearly apparent even at this level of analysis.

To more closely examine the specificity of our CR:tdTomato labeling approach in cortical regions, we performed immunolabeling for CR cell markers p73 and reelin in brain tissue sections from CR:tdTomato animals at P15 (**Fig 3.A**). We found that almost all cortical tdTomato+ neurons expressed the CR cell markers p73 and reelin, with very few neurons positive for both CR cell markers but not tdTomato (**Fig 3.B**). We performed cell counting of p73+, reelin+, and tdTomato+ neurons, and in total we quantified six different categories of cells, depending on the overlapped marker expression, from 11 different brain areas (**Fig 3. C**). Importantly, almost no cells expressed tdTomato without expressing at least one of the CR cell markers. The number of neurons expressing both CR cell markers but not tdTomato was consistently low throughout all the areas examined. Overall, for all areas counted, >95% of neurons expressing tdTomato co-expressed p73 and reelin (**Fig 3. D**), while the p73+/reelin+, tdTomato-group only amounted to ∼1.2% of all counted cells. To a similar degree, out of all identified CR cells (p73+/reelin+), >95% expressed tdTomato. These data demonstrate that our CR:tdTomato intersectional labeling strategy captures p73+ CR cells with excellent specificity and efficiency throughout different cortical regions.

**Fig. 3.**
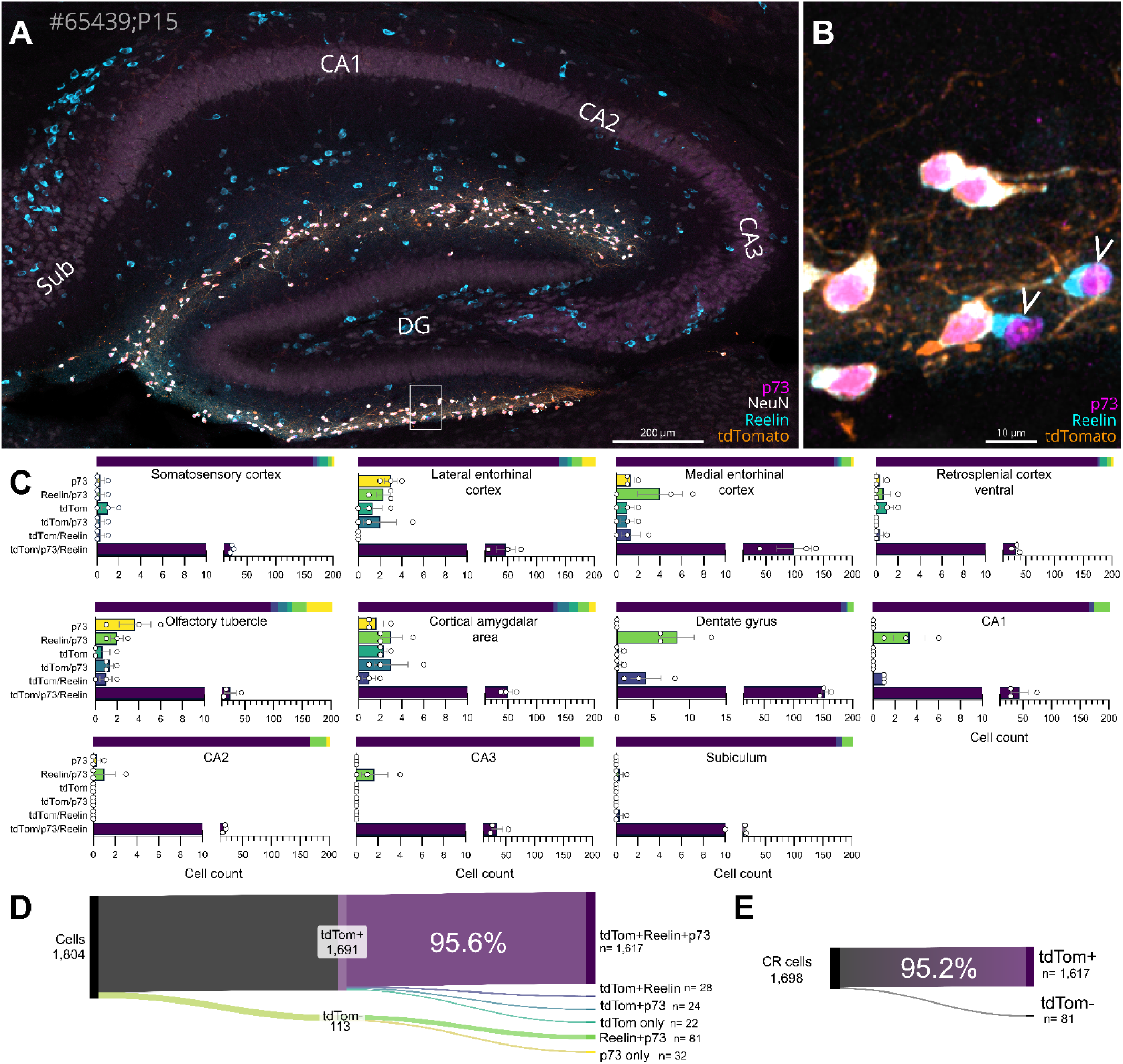
Quantification of CR cell specific marker expression in CR:tdTomato cells. (A) Sagittal section of the CR:tdTomato mouse hippocampus at P15, immunolabeled for tdTomato, p73, reelin, and NeuN. Scalebar, bottom right. (B) Zoomed in area from inset in A. White arrows point to neurons staining positive for p73 and reelin, but that do not express tdTomato. Scalebar, bottom right. (C) Bar plot for cell counts of six distinct marker overlaps from immunolabeled brain sections. One bar plot is shown from each area included in the quantification. Note the split in the X axes (cell counts) of the graphs, where after 10, the minor and major ticks are changed from one single to 10 for minor and 50 for major. (D) Sankey diagram showing summary of the counted cells categorized in distinct groups, being first split between tdTomato+ and tdTomato-. (E) Sankey diagram showing number of the putative CR cells that also express tdTomato.

### Expression and density of CR cells across areas and ages

Next, we set out to compare and quantify the number of CR cells across different brain areas and developmental timepoints. We first trained a segmentation algorithm using Ilastik (Berg et al., 2019), on a subset of the dataset, to facilitate automatic cell counting. Next, we aligned all sections to their corresponding Allen brain atlas image (Wang et al., 2020), using QuickNII (Puchades et al., 2019) and VisuAlign (Puchades et al., 2019). The segmentation and atlas alignments were merged using Nutil (Groeneboom et al., 2020) to quantify the number of tdTomato+ cells in all brain areas, which we then divided over the calculated layer 1 area volume to acquire a density value. All brain areas and the number of subjects used are listed in **Supp.Table 1**. By taking the normalized density values and assigning a color value distributed over a colormap, we created a heatmap of CR cell density (**Fig 4. A**). The heatmap illustrates various key features of CR cell density and distribution across age. At P1, sensory-motor areas exhibit a high density of CR cells, only surpassed by the hippocampus proper, dentate gyrus, and parasubiculum (PaS). However, the CR cell density in sensory-motor areas sharply decreases already by P15. In comparison, areas such as prefrontal (e.g., orbitofrontal, secondary motor area, infralimbic) and association cortices (e.g., retrosplenial, entorhinal, and perirhinal) have a lower density at P1, but show a relatively protracted persistence of CR cells through postnatal development (**Fig 4. A**).

**Fig 4.**
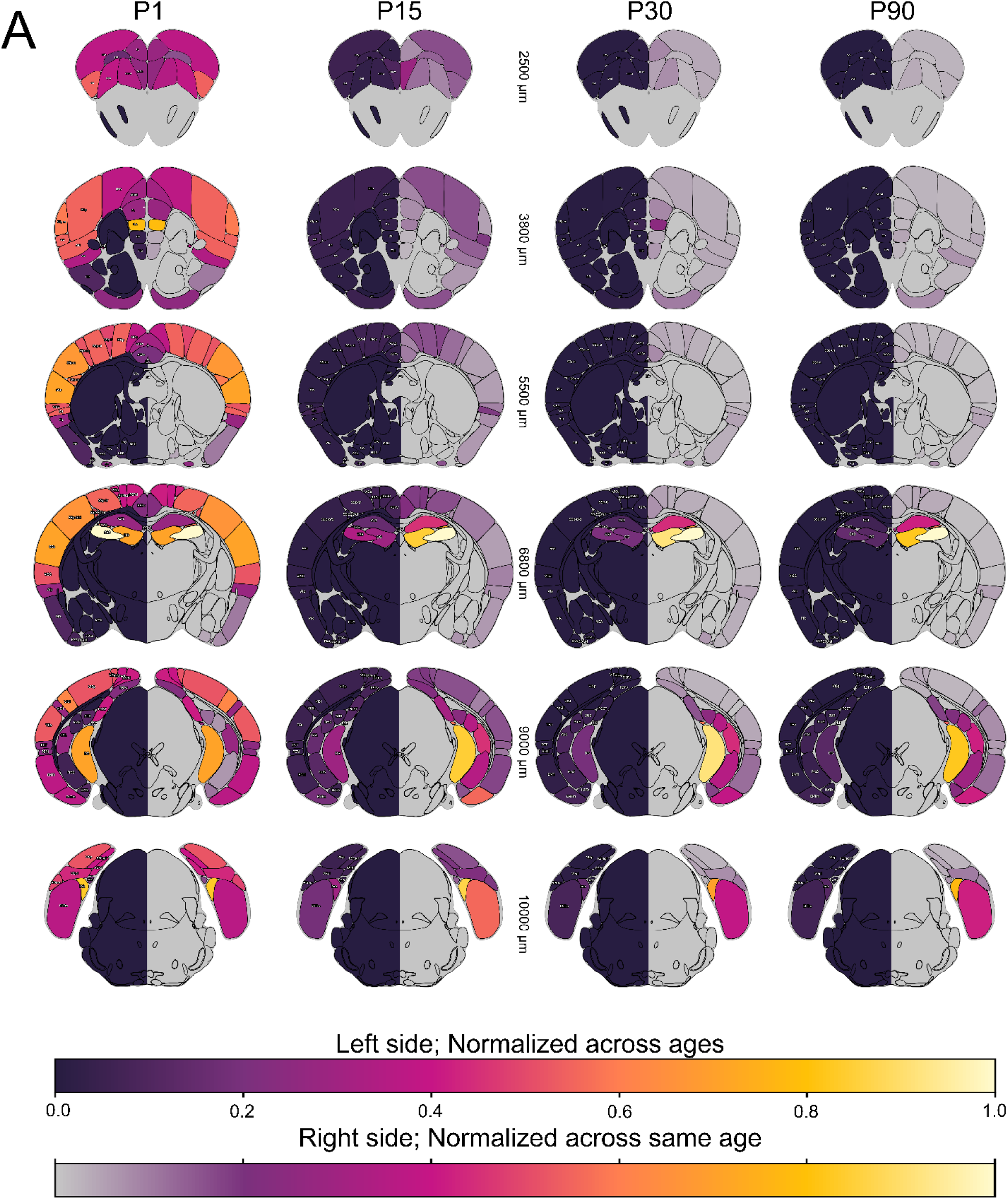
Heatmap for CR cell density across all relevant brain areas and ages. (A) Coronal section illustrations displaying CR cell density as heatmaps. For each brain section illustration, the left side shows a density value normalized across all ages. Right side, normalized across only same age. Left to right, P1 to P90. Top to bottom, increasing anterior-posterior (AP) positions. AP position stated across midline in µm distance from anterior apex of brain. Color bars indicate correspondence between color and density values. Top bar, for left side of each brain section illustration. Bottom bar, for right side of each brain section illustration. Abbreviated names for each cortical area are from the Allen Brain Atlas standard.

To better understand changes in density in CR cells across development, we decided first to focus our attention on four main cortical areas: the primary somatosensory cortex (SSp), which has been extensively studied with regard to CR cells (Damilou et al., 2024; Del Río et al., 1996; Genescu et al., 2022; Riva et al., 2019; Supèr et al., 2000), the hippocampus, and the two subdivisions of the EC, the medial (MEC) and lateral (LEC), as these are the major input regions to the hippocampus (Canto et al., 2008; Cappaert et al., 2015; Nilssen et al., 2019; Witter et al., 2017). CR cells are indeed most numerous at P1, in all the areas considered. At P1, CR cell density in the EC is comparable to that in other association cortices, but is lower than that observed in sensory-motor areas. The first obvious difference in CR cell trajectory starts appearing at P15, when the SSp displays a drastic decrease in CR cell number, compared to hippocampus. (**Fig 5.A**). Unexpectedly, MEC and LEC showed clear differences in CR cell persistence. As the animal ages, CR cell density decreases at a slower rate in MEC than in LEC. While MEC CR cell density follows more closely the rate of decrease of hippocampal CR cell density, LEC CR cell density decreases much more rapidly, a phenomenon evident at all ages after P1. At P30, and especially by P90, the CR cell density in MEC is still relatively high, as previously described only for hippocampus, while in almost all other cortical areas, CR cells are virtually gone. (**Fig 5.B**). Overall, across the analyzed timepoints, CR cells in the hippocampus exhibit a slower, stepwise decrease in density, while in the SSp, CR cell density rapidly decreases from P1 to P15. The change in CR cell density for MEC more closely resembles that of the hippocampus while the trajectory in LEC resembles that of the SSp.

**Fig. 5.**
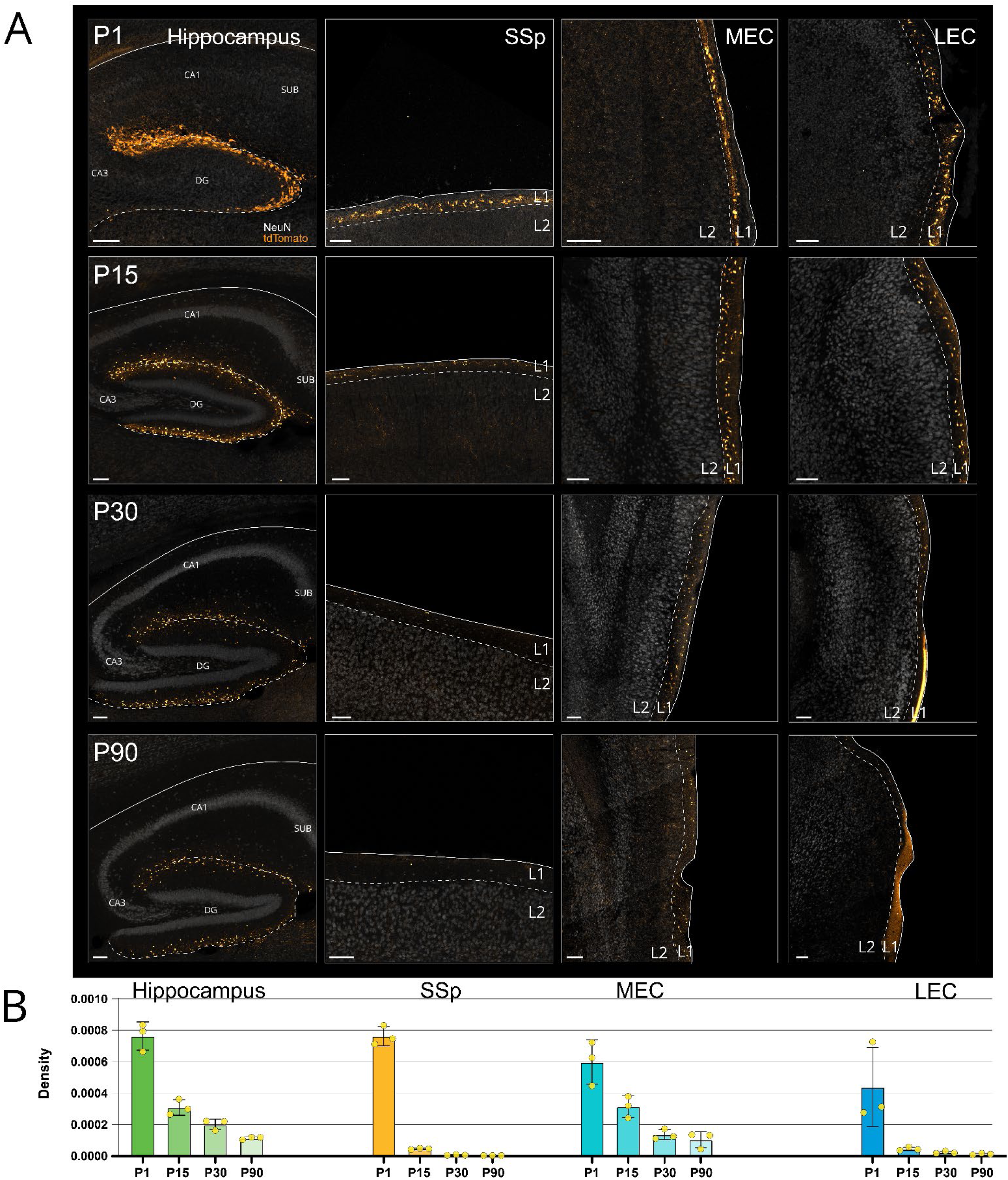
Selected brain areas illustrating differences in CR cell density trajectory over development. (A) Regions of interest from (left to right) hippocampus (dentate gyrus, CA1-3), primary somatosensory cortex (SSp), medial entorhinal cortex (MEC), and lateral entorhinal cortex (LEC). Top to bottom, P1 to P90. Dotted line delineates the hippocampal fissure or border between cortical L1 and L2. Scalebar for all images, bottom left: 100 µm. (B) Bar graphs for average CR cell density, from P1 to P90 across areas. Density calculated by dividing cell number over pixel volume for layer 1 in the respective areas. Dots indicate density from one animal. Error (SD) in black.

### Areal differences in the rate of CR cell loss across development

By analyzing CR cell density changes across ages throughout the whole brain, we uncovered that CR cells undergo cell death with distinct trajectories in different brain regions. To examine this phenomenon in more detail, we examined CR cell densities in a number of cortical regions across different levels of cortical hierarchy, in particular MEC and LEC, primary and secondary sensory cortices, association cortices, prefrontal cortex (PFC), and hippocampus. To be able to compare the time course of cell death across multiple regions, we assigned the density number of each region at P1 as 100%, with an arbitrary time point after P90 set as 0%, and then normalized each time point with these two extremes. When plotted, these time courses display the rate of disappearance of CR cells in the given area. As expected, the hippocampus exhibited the greatest postnatal persistence of CR cells compared to almost all other areas, with the exception of MEC, where remarkably, decay rates were even slower than that of the hippocampus (**Fig 6. A**). Together, these two regions exhibited the slowest decay in CR cell density, with both regions reaching a similar value at P30, and then similarly decreasing in density up to P90. The group with the most rapid CR cell decay rate is comprised of primary and secondary sensory cortices, losing most CR cells between P1 and P15, consistent with previous studies. In these regions, CR cell density is under 10% of P1 levels already by P15, and eventually reaches ∼0% at P30 (**Fig 6. A**). The association cortices, PFC and, LEC show a pattern of CR cell loss that is intermediate between the hippocampus and MEC, and the primary and secondary sensory cortices (**Fig 6. A**). Eventually, by P90, association cortices and PFC have lost all of their CR cells, like the sensory cortices, while LEC still remains at over 5%.

**Fig 6.**
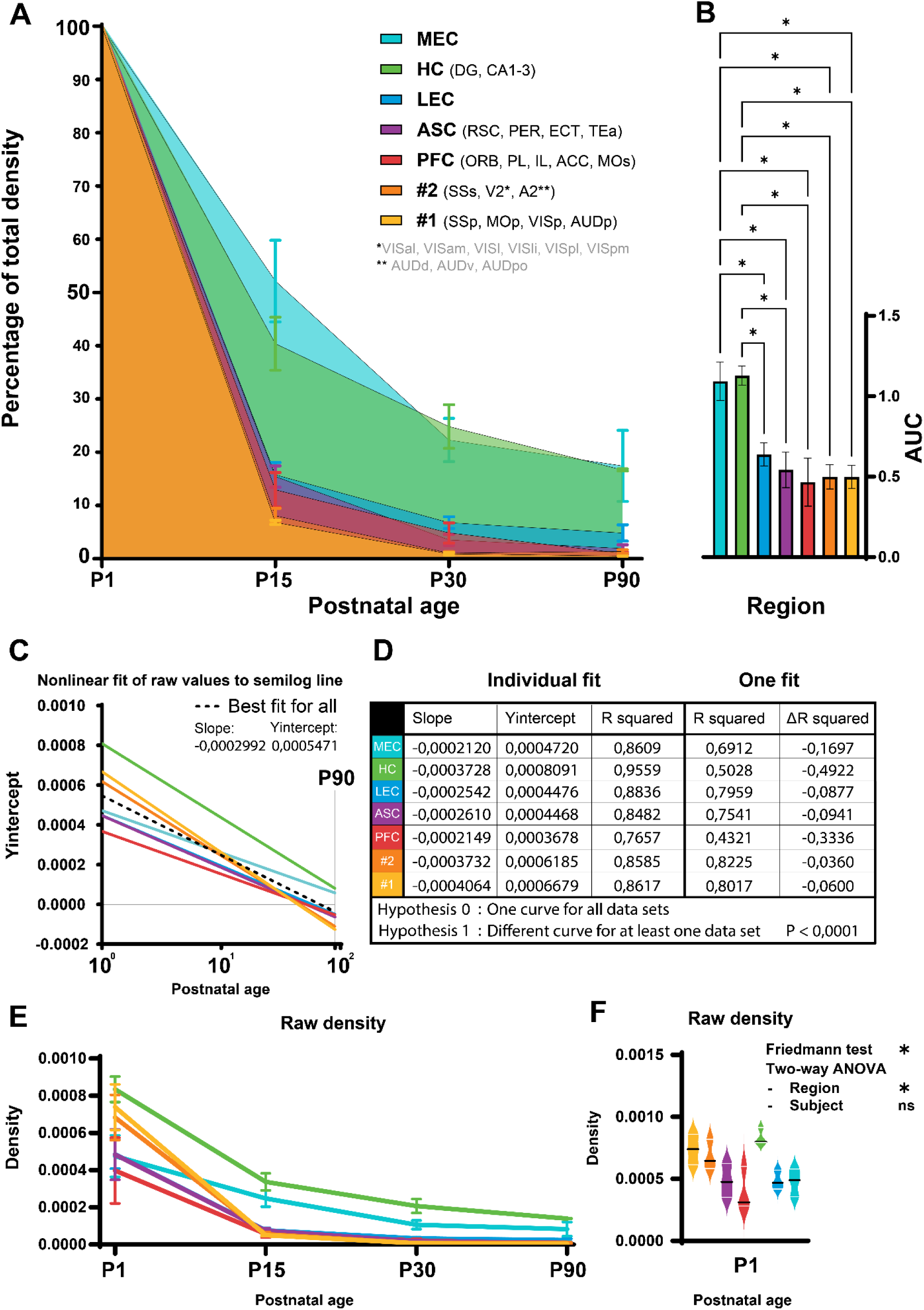
Differences in cell density decay across selected regions. (A) Line graph showing rate of decay of CR cell density for specific brain regions or areas. Regions and areas listed to the right. All regions start at 100%, which corresponds to the average density of the region at P1. 0% is artificially set after P90. (B) Bar plot showing area under the curve (AUC) for all regions. Brown-Forsythe ANOVA test, asterisk indicates P < 0.0001. No significant difference was discovered between MEC and HC. (C) Graph of slopes from nonlinear fit of raw values to semilog line, X-axis in log. (D) Table showing all slope and Yintercept values used to calculate the respective individual fit lines. R squared shows goodness of fit to the actual data. This is compared to R squared of a one fit line, with ΔR squared showing difference in fit between the individual fit and one fit. (E) Line graph for raw density values from all regions, highlighting difference in non-normalized CR cell density between regions. (F) Violin plot for raw density values from only P1. White lines show quartiles, black lines show median. Asterisk indicates P < 0.05.

In order to better analyze differences in the density decay of distinct areas, we calculated the area under the curve (AUC) for all the plotted regions and compared them with a Brown-Forsythe ANOVA test (**F\***(**DFn**, **DFd**) = 25.90 (6.000, 9.801), **P** <0.0001). A multiple comparison post-hoc test was then conducted to identify which regions had significantly different AUCs (**Fig 6. B**). To further compare the rate of decay between regions, we fit the raw density values from the regions across ages, using a nonlinear regression with a semi-log line (**Fig 6. C**). When comparing the individual fits to a single fit for all, it was revealed that at least one curve is different from the rest (**P**<0.0001), showing a notable decrease in goodness of fit (R squared) for several of the regions (**Fig 6. D**).

To perform this analysis and be able to compare different areas, we had to normalize the density value of CR cells for all brain regions. However, a different starting density of CR cells at P1 could affect our analysis of the decay rate. To understand if the number of CR cells at P1 was similar across the different regions of interest, we plotted the raw density values across age (**Fig 6. E**). A nonparametric Friedmann test indicated that the mean ranks of CR cell density at P1 were indeed different (**Chi-square** = 15.14, **P** = 0.0192), but a Dunn’s multiple comparisons test did not discover any significant differences when testing all possible pairs of regions. Lastly, we conducted a two-way ANOVA to discern if animal variability or the brain region were more likely to cause the variation in density. Interestingly, inter-subject variation could only explain 8.40% of the total variance (**F** = 2.37, **DFn** = 2, **DFd** = 12, **P** = 0.1353), whereas brain region could explain 70.36% of the total variance (**F** = 6.63, **DFn** = 6, **DFd** = 12, **P** = 0.0028 (**Fig 6. F**). While these data suggest that the starting density of CR cells is significantly different among the analyzed regions, it does not fully explain the differences in decay rates, but rather underscores them. This is particularly evident when looking at the MEC, a region with a very low decay rate, that is characterized by a lower density at P1 but that maintains a relatively high density at P90 when compared to other brain regions.

In order to understand what regional features or functions could contribute to persistence of CR cell density, we examined several additional neocortical areas (**Supp. Fig 2**). We therefore selected the primary visual cortex, known for its relatively delayed critical period (Fagiolini et al., 1994; Gordon & Stryker, 1996), and the barrel cortex, which matures earlier (Erzurumlu & Gaspar, 2012). Both primary visual cortex and barrel cortex (**Supp. Fig 2. C-D**) exhibited a CR cell density comparable to that of primary and secondary sensory-motor areas (**Fig 6. A-B**) with an AUC of ∼0.5. These results indicate that the timing of critical period onset is not a major contributing factor to the persistence of CR cells. However, primary visual cortex has a higher density of CR cells at P15 compared to the barrel cortex (**Supp. Fig 2. C**), which could be related to the later critical window in primary visual cortex and its relation to the timing of eye opening (Erzurumlu & Gaspar, 2012; Fagiolini et al., 1994). This data seems also to correlate with what we observed for PFC, which are known to have a protracted postnatal maturation (Chini & Hanganu-Opatz, 2021). Additionally, we chose to further analyze the retrosplenial cortex (RSC) in light of its interconnectivity with hippocampus and EC, and its importance in spatial processing and memory (Mitchell et al., 2018). Interestingly, the ventral portion of RSC (area 29; RSCv), which is highly interconnected with the hippocampal formation, displayed a higher persistence of CR cells at several postnatal ages, but especially at P15, compared to all other areas except for hippocampus and MEC (**Supp. Fig 2. C**). Out of all areas examined it was the only area to not have a significantly smaller AUC than hippocampus and MEC (**Supp. Fig 2. C-D**).

Overall, our analysis reveals that the decay rate of CR cells is region-specific, with sensory areas showing the fastest decay and the hippocampus and MEC having the slowest decay rate and greatest persistence of CR cells, even up to P90.

## DISCUSSION

This work presents a novel intersectional genetic labeling strategy for CR cells that exhibits a high degree of specificity and efficiency in isocortical and hippocampal regions. Use of this mouse cross (CR:tdTomato) allows for widespread and selective labeling of CR cells, which we took advantage of to investigate the distribution, density, and persistence of CR cells at four different developmental timepoints. Our results uncover nuances in the time course of CR cell disappearance across cortical and hippocampal areas, raising interesting questions on what regulates their survival. We show that (1) the CR:tdTomato mouse line is able to capture >95% of all p73+ CR cells in selected areas, and that >95% of all isocortical and hippocampal neurons expressing tdTomato are CR cells. (2) The expression of tdTomato in CR cells is stable throughout postnatal development, but at older ages, some non-CR neurons in the OB, cerebellum, deep layer piriform cortex, and dorsal medullar nuclei start to express tdTomato as well. (3) CR cell density and rate of disappearance exhibit substantial variation across different cortical areas during development. Our mapping revealed that the EC, association cortices, and PFC display a lower starting density of CR cells, but that these cells persist for much longer, even to early adulthood, in comparison to sensorimotor cortices. Interestingly, CR cell persistence in the MEC is comparable to that of hippocampus, whereas in the LEC it is more in line with that of association- and prefrontal cortices.

### Intersectional genetic strategy for improved Cajal-Retzius cell targeting specificity

Our intersectional genetic strategy offers several advantages over other mouse lines commonly used for CR cell manipulation (Pde1c-Cre: (Osheroff & Hatten, 2009); ΔNp73Cre: (Tissir et al., 2009; Yoshida et al., 2006); Frizzled10-CreERT2 (Gu et al., 2009); CXCR4-EGFP (Marchionni et al., 2010). The most pertinent comparison is to the single transgenic Pde1c-Cre^+/−^mouse line. Although successfully employed previously to manipulate CR cells, most recently by our own group (Glærum et al., 2024), it does show transgene expression in other isocortical neuronal populations (Osheroff & Hatten, 2009). This could potentially pose a problem when trying to manipulate the cells, both when crossing it with specific floxed mouse lines and when injecting viral constructs directly into, or through, cortical areas, as the construct could get introduced in non-CR cell populations. The use of the NDNF-flox-FlpO knock in line in combination with Pde1c-Cre offers improved access to almost all p73+ CR cells, with much less potential off-target expression. Furthermore, the design of the NDNF-flox-FlpO driver allows for local targeting of intersected cell populations with Flp-dependent (fDIO) AAV constructs since only those NDNF+ cells that express Cre (i.e., Pde1c-Cre in our case) express FlpO.

Another commonly used mouse line is the ΔNp73Cre line (Tissir et al., 2009), which specifically expresses Cre recombinase in all p73+ neurons. However, p73 is expressed by other populations besides CR cells, including neurons in the preoptic area, gonadotropin-releasing hormone expressing neurons, and cells in the vomeronasal organ and choroid plexus (Tissir et al., 2009; Yang et al., 2000). More importantly, when expressing diphtheria toxin A under the control of Cre, Tissir and colleagues (Tissir et al., 2009) noted that only 72% of hippocampal CR cells were ablated, suggesting a significant population of CR cells cannot be targeted with this mouse line. When using the Wnt3a-Cre mouse line (Yoshida et al., 2006), the authors achieved a reduction of only 35% in CR cell number, while a combination of both lines yielded a CR cell number reduction of 84%. In addition, the Wnt3a-Cre mouse line is rather unspecific at postnatal ages, as many granular cells in the dentate gyrus express Wnt3a (Pinnock et al., 2010).

Our quantification of tdTomato expression in our CR:tdTomato mice indicated that >95% of CR cells were labeled, demonstrating that this mouse line is to date the most specific and effective genetic approach to target CR cells.

### Persistence of Cajal-Retzius cells varies across distinct brain regions

The difference in time course of cell death for CR cells between isocortex and hippocampus has been a recognized feature of CR cells for several decades. Still, a nuanced examination of this time course over finer parcellated brain areas has been lacking. By extending our analyses to more brain areas, we have been able to appreciate these nuances in a new way. Based on the results from the density heatmaps, we chose to more closely examine the density of CR cells over time for three distinct groups of regions: the EC and hippocampus, association cortices and PFC, and primary and secondary sensorimotor cortices. These three groups are partly segregated in the CR cell persistence graph (**Fig 6. A**). As expected, primary and secondary sensorimotor cortices and the hippocampus show a very different CR cell density trajectory across development, with CR cells persisting longer in the hippocampus. Intriguingly, association cortices and PFC exhibit an intermediate degree of CR cell survival compared to hippocampus and sensory cortices. What might account for the prolonged survival of CR cells in hippocampus and specific cortical areas? We speculate that this phenomenon relates to the differences in the maturation trajectories and functions of these brain regions.

The PFC is famously characterized as a hub for many cognitive and social abilities that emerge relatively late during postnatal maturation (Chini & Hanganu-Opatz, 2021). Indeed, the PFC is noted to have a protracted developmental timecourse compared to other isocortical regions (Chini & Hanganu-Opatz, 2021; Huttenlocher & Dabholkar, 1997; Selemon, 2013), which in turn is reflected in the delayed maturation of synaptic stabilization in PFC circuitry (Chini & Hanganu-Opatz, 2021; Kroon et al., 2019; Petanjek et al., 2011; Elston et al., 2009; Hoops et al., 2018). In our data, the PFC only showed a small difference in CR cell trajectory compared to other isocortical areas. However, several regions of PFC are positioned along the midline, which proved difficult to quantify in sagittal sections, becoming clearer in coronal sections (**Supp. Fig 2. A-B**). Thus, it is possible our density analysis underestimates the number of CR cells in the PFC, as two thirds of the dataset used in this study were collected from sagittal sections. Observation of CR cell persistence in these areas in coronal sections suggested a somewhat slower CR cell decay compared to primary visual cortex and the barrel cortex, supporting the idea that CR cell persistence could be partly explained by the local maturational timeline. Future experiments aiming at manipulating the maturation of different regions while assessing the density of CR cells could help in clarifying this point.

### Timing of Cajal-Retzius cell persistence is markedly different between medial- and lateral entorhinal cortex

Within the subregions of the EC, we observed a striking difference in how long CR cells persisted through development. Even though the two subdivisions of the EC have a similar cytoarchitecture and output relationship to the hippocampus, they have distinct differences in what inputs they receive, and each harbor distinct functional cell types (Nilssen et al., 2019; Witter et al., 2017). For example, the main output neuronal types in layer 2 are morphologically distinct in MEC and LEC, being comprised of stellate cells and fan cells, respectively (Canto & Witter, 2012a, 2012b).

One possible explanation for different CR cell persistence between the two EC subdivisions could be related to the embryonic development of this region. First, based on ^3^H-thymidine autoradiography, Bayer (Bayer, 1980) reported that cells in the EC are generated in the order of lateral, to intermediate, to medial. The earlier timecourse of neurogenesis of LEC principal neurons compared to MEC could be related to the earlier disappearance of CR cells in the former. Second, MEC and LEC have been suggested to originate from different pallial regions (Abellán et al., 2014; Medina et al., 2017). MEC shares its origin, the medial pallium, with all other hippocampal structures, meaning not only the hippocampus, but also the presubiculum (PrS) and PaS, which also have a relatively high CR cell density at older ages (**Fig. 4**). On the other hand, LEC is thought to originate from a more dorsolateral part of the pallium, similar to other associative areas like the perirhinal cortex (PER) and orbitofrontal cortex (Abellán et al., 2014; Medina et al., 2017; Nilssen et al., 2019). It is therefore an intriguing possibility that the observed differences in CR cells persistence among regions might be related to the developmental origin or timeline of the region in which they are located.

Another factor that could influence CR cell persistence is the difference in the invasion of the input fibers in layer 1 between the EC divisions. LEC is proposed to integrate an abundance of associative signals, with major inputs from PER and postrhinal cortex (Doan et al., 2019; Nilssen et al., 2019). Additionally, olfactory and piriform signals strongly innervate LEC (Burwell & Amaral, 1998; Nilssen et al., 2019). Fibers from the MOB reach LEC already at P3-P4 (Gretenkord et al., 2019), and have been shown to influence LEC development, for example by driving dendritic growth (Chen et al., 2023). The fact that smell is one of the first senses to develop, affecting behavior already in utero (Stickrod et al., 1982) could lead to an earlier maturation of the LEC layer 1 circuit, compared to MEC. MEC receives primarily input from PrS and PaS, which terminate in all four principal layers of MEC (Canto et al., 2012), and RSC, which terminate in deep layers (Doan et al., 2019; Nilssen et al., 2019; Sugar & Witter, 2016). PrS and PaS fibers arrive in MEC between P9 and P10 and continue developing until P28-P30 (Canto et al., 2019). Functionally, the MEC also seems to mature later than the LEC. For example, grid cells only fully attain their mature properties at around four weeks of age in rats (Langston et al., 2010), being reliant on the animal’s experience in exploring its environment (Loewen et al., 2005; Ulsaker-Janke et al., 2023). Thus, the different input, and the timing of when functionality emerges in the two EC divisions might explain at least in part why CR cells persist longer in MEC compared to LEC.

An intriguing comparison can be made between the difference observed in the CR cell density trajectory in MEC compared to LEC, and the difference between RSCv compared to dorsal RSC (A30; RSCd). Both subregions of RSC have been suggested to be involved in spatial memory, but they are interconnected with different regions (Mitchell et al., 2018). As reported here, RSCv has a prolonged persistence of CR cells compared to other neocortical areas, whereas dorsal RSCd shows an earlier decay of density and smaller AUC. RSC likely processes allocentric to egocentric transformations of navigational signals, but RSCd, receiving mainly visual inputs, has been found to be more biased towards egocentric information in comparison to RSCv, which is more interconnected with the hippocampal formation (Aggleton et al., 2021; Alexander et al., 2020). MEC and hippocampus are thought to predominately support allocentric spatial processing (Keefe & Nadel, 1978), while LEC is proposed to process more egocentric signals (Wang et al., 2020). Thus, it is interesting to speculate that CR cell persistence might be related to the nature of spatial information processing in each area.

### Correlation between Cajal-Retzius cell origin and persistence

One interesting possibility is that differences in CR cell decay rate could be due to the distinct embryonic origins of these cells. To date, four different embryonic regions have been identified to give rise to CR cells (Causeret et al., 2021). CR cells of different origin exhibit similar subpial positioning and morphology, as well as the expression of reelin, but express different molecular markers and populate different parts of the developing neuroepithelium (Causeret et al., 2021). For example, CR cells with a medial origin, coming from the hem, septum, or thalamic eminence (TE) express the marker p73, while CR cells from the PSB are p73-negative (Griveau et al., 2010; Ruiz-Reig et al., 2017). Given that in our CR:tdTomato mouse, over 95% of the CR cells express p73, as analyzed at P15, it is possible that we are capturing a very small population of PSB derived CR cells, as our characterization showed a minor number of reelin+, tdTomato+, p73-cells in all areas characterized (**Fig 3. C**)

CR cells from the hem distribute to the caudomedial and dorsal cortical regions, including hippocampus, while septal CR cells distribute medially and dorsoventrally over the more rostral part of the brain. PSB CR cells distribute more ventrally along the rostrocaudal axis, while CR cells from the TE are distributed caudoventrally (Bielle et al., 2005; Ruiz-Reig et al., 2017; Takiguchi-Hayashi et al., 2004). However, the distribution of CR cells from different origin is not homogenous across cortical areas, with some areas having a mixed pool of CR cells from several origins. For example, the piriform cortex likely has CR cells arising from the PSB, septum, and TE (Bielle et al., 2005; Causeret et al., 2021; Ruiz-Reig et al., 2017). As CR cells persist for the longest in hippocampus, EC, and along the midline (RSC, PFC), both the hem- and septal derived CR cells could persist into late development. Still, the hippocampal populations, which persist for longest, have mostly hem origin, as they express Wnt3a during development (Lee et al., 2000; Louvi et al., 2007). The origin of CR cells in the MEC cells has not been thoroughly assessed, but Quattrocolo and Maccaferri (Quattrocolo & Maccaferri, 2014) reported CR cells labeled by the Wnt3a-Cre mouse line in the MEC in the first postnatal month. It is therefore possible that a substantial population of MEC CR cells originates from the cortical hem, suggesting their decay rate could be related to their origin. Future analysis of the embryonic origin of CR cells in different cortical and hippocampal region will help answer this question.

## CONCLUSION

CR cells are essential cells for the correct organization of cortical layers and establishment of early prenatal brain circuitry. The postnatal role of these cells is just beginning to be appreciated to a fuller extent now that we are developing more advanced tools to observe and manipulate them. Here, we have developed a novel genetic labeling approach (CR:tdTomato) which accurately and efficiently targets CR cells throughout the whole mouse brain. Using this strategy, we were able to uncover both the distribution and decay rate of CR cells over the entire brain. Our results reveal that CR cell density is not uniform throughout postnatal development, and that the rate of their disappearance varies significantly between different brain regions. CR cells persist longer in association cortices, PFC, EC, and hippocampus. A difference in CR cell decay rate within the EC also highlights possible developmental and maturational differences between MEC and LEC, which might be related to their origin and functional differences. We have contemplated several different causes for these observed differences in CR cell decay rate, but future studies are required to resolve the underlying mechanism of CR cell persistence. Likely, CR cell persistence is governed by a multitude of factors relating to their postnatal function and the maturation trajectory of the specific area where they reside. Further studies of the postnatal function of CR cells combined with lineage information of where the particular CR cells originate from will help us elucidate how these unique cells support specialized postnatal development programs in different brain areas.

## Acknowledgements

We would like to thank B.A. Zaharia for excellent technical assistance; H.M. Møllergård for genotyping; P.J.B. Girão for assistance with microscopy; K.Dunville for skillful statistical assistance; M.J.Nigro, and all the members of the Quattrocolo lab for their insightful comments throughout this project. We further thank the staff in the animal facility at the Kavli Institute for Systems Neuroscience.

The work was supported by a Research Council of Norway (RCN) FRIPRO grants to G.Q. (grant number 324305), RCN Centre of Excellence grants (Centre for Algorithms in the Cortex, grant number 332640; Centre of Neural Computation, grant number 223262; to G.Q.) and the Kavli Foundation (to G.Q.). The experiments were performed at the NORBRAIN Facility, Norwegian University of Science and Technology (NTNU) (grant number 295721).

## Author contribution

Conceptualization, K.M and G.Q.; supervision, G.Q.; experiment design, K.M., and G.Q.; resources, all authors; data collection, K.M., visualization, K.M.; writing, original draft, review and editing, discussion and comments, all authors; funding acquisition: G.Q.

## Supplementary figures

**Fig 1S.**
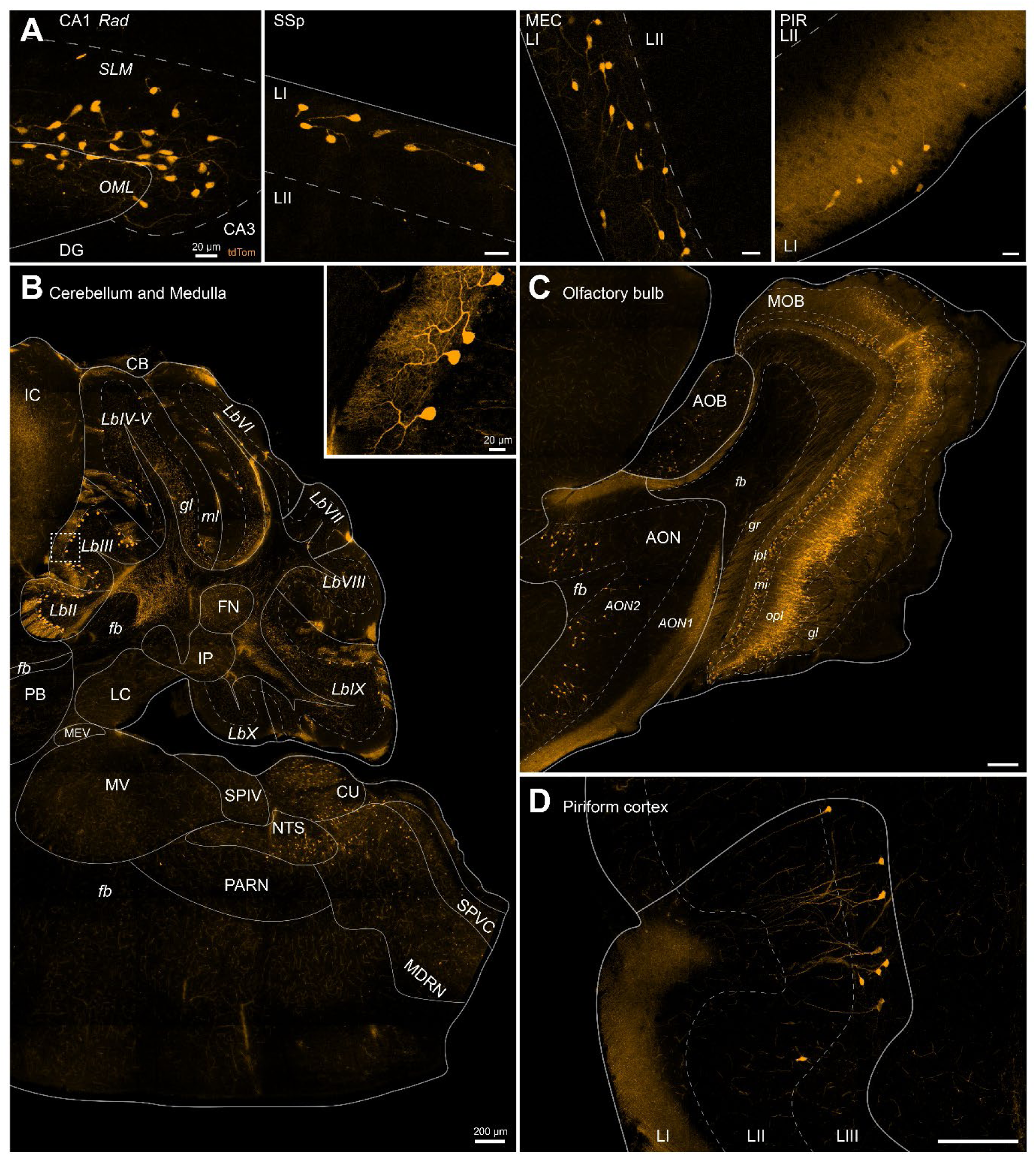
tdTomato-positive cells labeled in the CR:tdTomato mouse line. (A) Four representative images of CR cells from different areas, displaying CR cell morphology. From left to right: Hippocampus, primary somatosensory cortex, medial entorhinal cortex, piriform cortex. Scalebar, bottom right, 20 µm, same for all panels (B) Cells in cerebellum and medulla. Image in top right shows zoom from inset with dashed line. Scalebar, bottom right, 200µm, inset, 20 µm. Same for all following panels (C) Olfactory lobe (D) Layer III cells in piriform cortex. All delineations done manually over corresponding Allen brain atlas image, using NeuN (not shown) as counterstain. All abbreviations in Allen Brain Atlas standard.

**Fig 2S.**
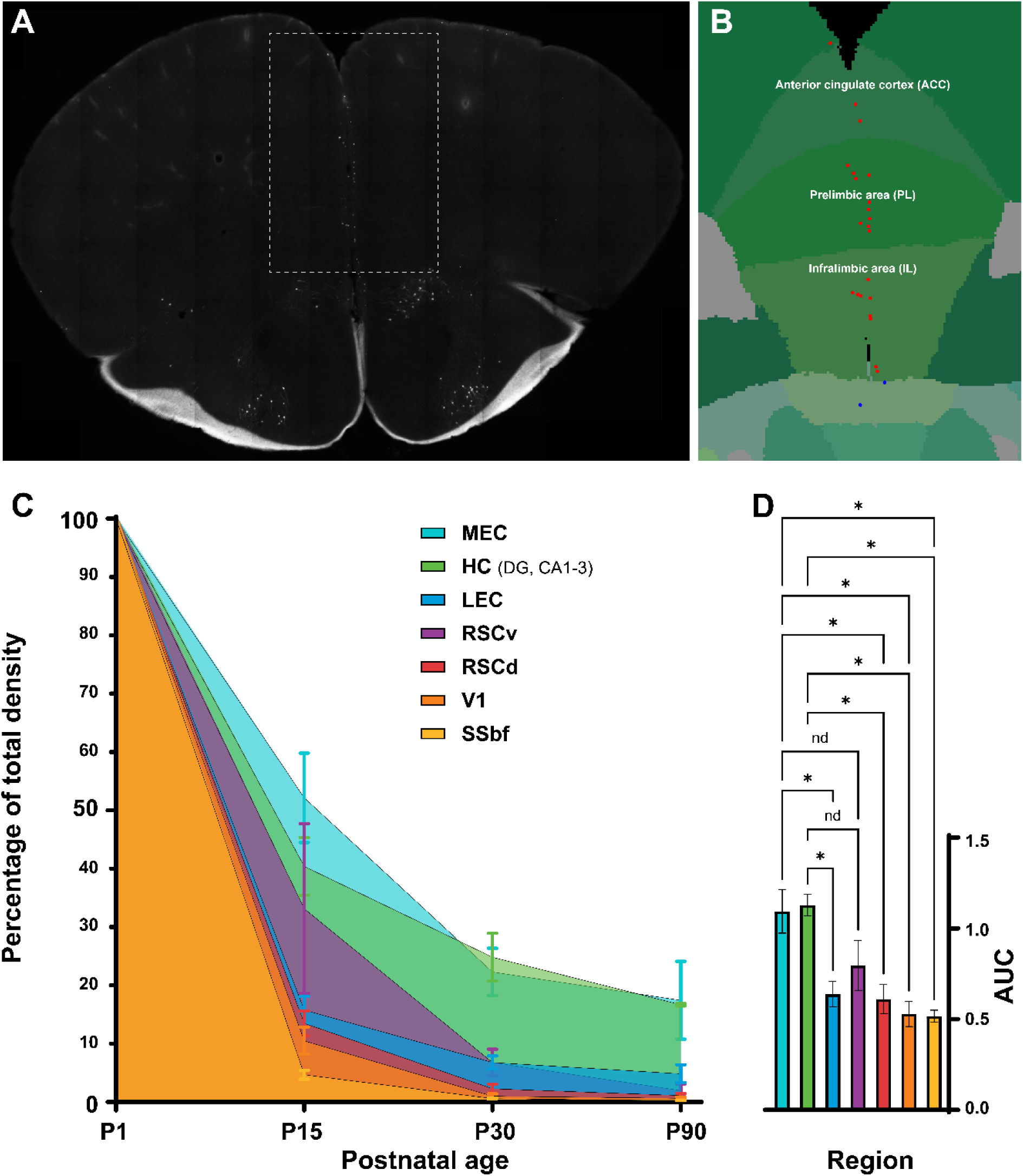
Differences in persistence across brain regions. (A) Widefield scanner images of a P30 coronal section from a CR:tdTomato mouse, stained for tdTomato, showing parts of PFC (Prelimbic, infralimbic, anterior cingulate cortex). (B) Atlas illustration with segmented tdTomato+ cells from automatic counter. Taken from inset in A. (C) Line graph showing density from respective areas across four ages. Areas normalized over only same area, to set start density at 100%. (D) Bar plot showing area under the curve (AUC) for all regions. Brown-Forsythe ANOVA test, asterisk indicates P < 0.0001.

## Supplementary Table

**Supp. Table 1.**
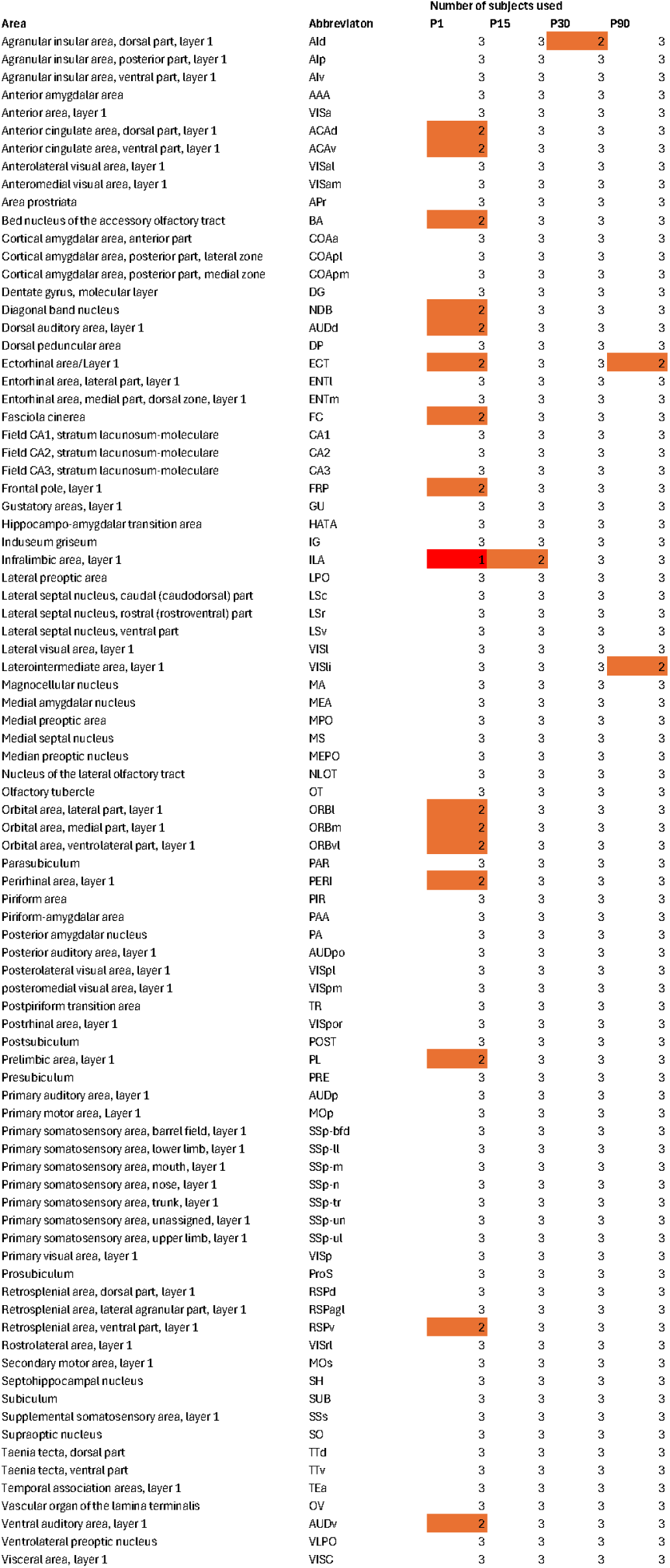
All brain areas with abbreviations used in counting. For all brain areas the number of subjects used for counting is listed under the respective age. Areas where we had to remove subject(s) are colored with orange (only two subjects used) or red (only 1 subject used). All abbreviations in Allen Brain Atlas standard.

## Notes

### Competing Interest Statement

The authors have declared no competing interest.

